# Insights into Trypanosomiasis Transmission: Age, Infection Rates, and Bloodmeal Analysis of *Glossina fuscipes fuscipes* in N.W. Uganda

**DOI:** 10.1101/2023.11.21.568004

**Authors:** Lucas J. Cunningham, Johan Esterhuizen, John W Hargrove, Mike Lehane, Jennifer Lord, Jessica Lingley, TN Clement Mangwiro, Mercy Opiyo, Iñaki Tirados, Steve J. Torr

**Affiliations:** Department of Vector Biology, Liverpool School of Tropical Medicine; Agricultural Research Council - Onderstepoort Veterinary Research, Pretoria, South Africa; DSI-NRF Centre for Epidemiological Modelling and Analysis, University of Stellenbosch, Stellenbosch, South Africa; Bindura University of Science Education, Bindura, Zimbabwe; Malaria Elimination Initiative, Institute of Global Health Sciences, University of California San Francisco, San Francisco, CA, USA; Centro de Investigação em Saúde de Manhiça (CISM), Manhiça, Mozambique

## Abstract

**Background:** Tsetse flies (*Glossina*) transmit species of *Trypanosoma* which cause human African trypanosomiasis (HAT) and animal African trypanosomiasis (AAT). Measuring the infection status of wild-caught tsetse is an important part of operations to control HAT. In north-west Uganda, we conducted field studies over a 15-month period to compare classical, microscope-based, and PCR-based methods of detecting trypanosomes in tsetse. We also quantified the age structure and host preferences of tsetse.

**Methods:** Using Pyramidal traps placed along the Kochi River, in Koboko district, 12512 G. fuscipes fuscipes were caught. A subset of females (¬n= 5051) and males (n= 1221) underwent dissection wherein mouthparts, midguts and salivary glands were screened for trypanosomes. Additionally, the age of the females was estimated using ovarian ageing and the trypanosome-positive status of 1931 females and 438 males was investigated by ITS PCR. Further the bloodmeal sources of 131 tsetse were identified using vertebrate cytochrome b PCR.

**Results:** Infection rates estimated were significantly greater (1.9-9.3 times) using the PCR-based method compared to the classical dissection-based method. Positive rates for *T. brucei* sl, *T. congolense* and *T. vivax* were 1.6% (1.32-2.24), 2.4% (1.83-3.11and 2.0% (1.46-2.63), respectively by PCR. The abundance and age structure of tsetse populations were relatively stable and the slight seasonal four-fold variation in abundance appeared to be correlated with rainfall. Analyses of age structure suggests a low natural daily mortality of 1.75% (1.62-1.88). The majority of bloodmeals were identified as cattle (39%, 30.5-47.8) and human (37% of meals, 28.4-45.6).

**Conclusion:** PCR provides a more sensitive and specific method of estimating the infection rates of all pathogenic species of trypanosome circulating in Koboko. The seasonally stable abundance, low mortality rate and high proportion of bloodmeals from humans may explain, in part, why this district has historically been a focus of sleeping sickness.

**Author summary:** Tsetse flies (*Glossina*) transmit African trypanosomiasis in humans (‘sleeping sickness’) and livestock (‘nagana’). Identifying trypanosome species and quantifying infection rates in tsetse is important in understanding and controlling trypanosomiasis. Traditionally this relied on microscopic examination of tsetse, however, this method is unable to reliably identify different species of trypanosome. We compared microscopy and PCR, trypanosome-detection methods, using *G. fuscipes fuscipes* collected from traps deployed in Koboko district, in North-West Uganda.

Our results show that the PCR-based method was 1.9-9.3 times more sensitive than the classical approach and able to identify different species of *Trypanosoma*, including mixed infections. In collecting tsetse we also obtained data on the seasonal abundance, age and diet of the tsetse population. Analyses of these results showed that natural mortality was low (1.75%/day) among adult females and only a slight seasonal variation in abundance and age structure occurred. The most important hosts were cattle and humans, providing 39% and 37% of bloodmeals, respectively.

The PCR method provided a sensitive means of identifying trypanosomes to the species level. The longevity and diet of tsetse in Koboko may explain, why this district was a persistent focus of disease prior to the deployment of Tiny Targets to control tsetse.

## Background

Tsetse flies (*Glossina*) transmit species of *Trypanosoma* which cause human African trypanosomiasis (HAT) and animal African trypanosomiasis (AAT). Across Central and West Africa, riverine tsetse (e.g., *G. palpalis*, *G. fuscipes*) transmit *T. brucei gambiense* which causes Gambian HAT (gHAT), a chronic and anthroponotic form of the disease. In East and Southern Africa, savanna tsetse (e.g., *G. morsitans*, *G. pallidipes*) transmit *T. b. rhodesiense* which causes Rhodesian HAT (rHAT), an acute zoonosis. Both forms of HAT are fatal unless treated with drugs. Between 2000 and 2021, the annual number of cases of gHAT and rHAT reported globally by the World Health Organisation has declined by 98% (747/25841) and 86% (55/709) respectively [1].

Uganda is the only country with both forms of HAT and has experienced severe epidemics of both over the last century. The most recent epidemic occurred towards the end of the last century and numbers have declined markedly since then. During 1990-1999, for instance, the mean number of cases of gHAT and rHAT reported annually in Uganda were 1384 and 516 cases/year, respectively, compared to 5 and 25 cases/year, respectively, for the period 2012-2021 [1]. The decline in cases of gHAT has been achieved through a combination of mass screening and treatment of the human population [2], coupled with use of Tiny Targets to control tsetse [3, 4]. Control of rHAT cases has been achieved through mass treatment of cattle with trypanocides and insecticides [5, 6]; in Uganda, cattle are important reservoir hosts for *T. b. rhodesiense* [7] and a source of bloodmeals for tsetse [8]. Recognising the progress made against HAT, the WHO provided formal validation in 2022 that gHAT has been eliminated as a public health problem in Uganda [9].

Given that gHAT is now a relatively rare disease, new strategies are needed to ensure that incidence remains low. One important component of the new strategy is to monitor the distribution and abundance of tsetse populations [4, 10–12], and, less commonly, to analyse tsetse for the presence of pathogenic subspecies of *T. brucei* [12]. Classically, detection and identification of trypanosomes was based on microscopic examination of tsetse [13] but this approach faces three important limitations. First, a fly observed to have trypanosomes present in the salivary glands and midgut is presumed to be infected with *T. brucei*. However, this method is unable to distinguish between *T. brucei brucei*, which is not pathogenic to humans, and the pathogenic subspecies *T. b. gambiense* and *T. b. rhodesiense.* Second, the presence of trypanosomes in the midgut only may signify an immature infection with *T. brucei* or *T. congolense*. Third, mature co-infections of, say, *T. congolense* and *T. brucei* would be incorrectly identified as being an infection with *T. brucei* only. These limitations have been highlighted in empirical studies of savanna tsetse in Tanzania [14] and riverine tsetse in Côte d’Ivoire [15]. Following comparison of dissection and PCR-based methods, Lehane *et al*. [14] concluded that “Given that so much of our understanding of the epidemiology of trypanosomes in tsetse flies is based on the Lloyd & Johnson (1924) dissection technique these results are potentially alarming and the study needs to be repeated as a matter of urgency.”

Another problem is that a mature, salivary gland, infection of *T. brucei* takes at least 15 days to develop after a fly feeds on an infected host [16]. This, coupled with the fact that traps tend to be biased towards the capture of old flies [17] means that the infection rate among trap-caught flies will tend to be greater than in the population as a whole.

Recognition of the limitations of dissection-based methods has led to greater use of PCR as a diagnostic tool [18–20]. However, studies using PCR-based methods alone are unable to distinguish tsetse with mature infections from immature ones, and it is only the former that are infectious. Moreover, detection of trypanosome DNA does not necessarily indicate the presence of an active infection in tsetse [21].

Following the exhortation of Lehane *et al.* (see above) we aimed to test the hypotheses that, at least for *G. f. fuscipes*, dissection-based analyses of tsetse do not provide a reliable estimate of the of infection with *Trypanosoma*. To test this hypothesis, we compared the dissection- and PCR-based estimates of trypanosome infections in a tsetse population sampled over a 15 month period in northern Uganda. Conducting the study over a period of 15 months also allowed us to examine seasonal fluctuations in abundance, age structure and trypanosome prevalence. The study of DNA from large numbers of tsetse also allowed us to answer a secondary research question: What are the important hosts in the diet of the tsetse population? Taken together, the various sorts of data we produced are useful in the interpretation of sampling operations and the planning of control in N-W Uganda, and are of practical and theoretical interest elsewhere.

## Methods

### Study site

All field studies were conducted, between April 2013 and July 2014 along the Kochi River in the Koboko district of north-west Uganda. The area is predominantly agricultural with crops such as cassava, tobacco, millet and sesame being farmed. Potential hosts of tsetse include various livestock species (cattle, goats, sheep and pigs) and monitor lizards [22]. Koboko and the surrounding districts have long been a focus of sleeping sickness, with reports dating back to 1905 [23]. The primary vector of *T. b. gambiense* in this district is *G. f. fuscipes* [24], a riverine species of tsetse.

### Tsetse sampling

Pyramidal traps [25] were placed at four sites along the Kochi river, <1m from the river’s edge and >100 m from each other. These traps formed part of a network of traps deployed initially to monitor the impact of Tiny Targets on tsetse populations [24]. The traps along the Kochi River monitored tsetse numbers in an area where Tiny Targets were not deployed and hence provide data on a natural, uncontrolled, population of tsetse. Traps were checked twice daily, at ∼07:30 h and ∼15:30 h. Tsetse were brought back to the laboratory in a cool box to reduce mortality from desiccation and heat stress.

### Tsetse dissection

Male and female tsetse were dissected to determine the presence of trypanosomes [26]. Female flies were also subjected to ovarian dissection to determine their age. The dissections were carried out by trained technicians using fine forceps and Zeiss Stemi 2000 dissection microscopes. Salivary glands, midgut and mouthparts were dissected out and then examined at 200× and 400× magnification using a compound microscope fitted with a dark-field filter. Tissues (e.g., salivary glands, midgut, mouthparts) were identified as positive if live, i.e., mobile, trypanosomes were observed within them. Infections were assigned putatively to a species of *Trypanosoma* based on the tissue(s) where trypanosomes were found [26].

### Molecular identification of Trypanosoma

Following dissection and scoring for infection and age, the mouthparts, salivary glands and midgut were each stored individually in 60µl of absolute ethanol, for subsequent molecular analysis; tissues from flies were stored in this manner regardless of whether trypanosomes had been found by microscopic examination. The samples were stored in 96-well PCR plates, with each plate holding tissues from 32 flies. Of the 6869 tsetse dissected, a subsample of 2370 was selected for PCR-based analyses to identify trypanosomes; a further subsample of 768 flies was selected for analysis aimed at identifying the source(s) of their bloodmeals.

### DNA Extraction

DNA was extracted from the samples as follows. The plate was spun at ∼6,000 rpm for 30 seconds and then the lids from the plates were removed and the ethanol evaporated off in an oven at 56°C for 1-2 hours. After evaporation, 135µl of DNA extraction buffer was added. The buffer comprised 1xTE/5% Chelex/ 1% proteinase K (20mg/ml). To elute the DNA, samples were incubated at 56°C for an hour, followed by a second incubation step at 93°C for 30min. The supernatant (100µl) was then pipetted off into a new 96-well plate, being careful not to pick up any of the Chelex beads.

### PCR protocols

#### Trypanosoma

For each fly, 2µl of DNA eluate was taken from the three tissues and pooled together, effectively reconstituting the fly. This pooled sample was then screened with the multiplex ITS primers [27], to screen for *T. congolense*, *T. brucei* and *T. vivax.* Any positive fly then had eluates from the three tissues tested separately using the same primers.

#### Blood meal analysis

The DNA eluate from a subsample comprising 768 midguts was screened using sequencing primers, designed to amplify vertebrate DNA by targeting a ∼470bp region of the cytochrome b gene. This allowed the host DNA in the bloodmeal to be targeted and amplified [28]. The primer sequences are as follows; forward ‘TACCATGAGGACAAATATCATTCTG’, reverse ‘CCTCCTAGTTTGTTAGGGATTG ATCG’. These primers amplified a ∼470bp region of the vertebrate mitochondrial genome. Once amplified the PCR product was then cleaned using the QIAquick PCR purification kit (Qiagen, Manchester, UK) and sequenced by SourceBioScience (Nottingham, UK).

### Environmental data

To assist in the interpretation of temporal variation in the catches, age structure and infection rates of tsetse, we collected local estimates of monthly rainfall and mean temperature. These were obtained from the Global Precipitation Climatology Centre (GPCC) and HCN_CAMS Gridded 2m Temperature (Land) data, respectively. All data were provided by NOAA PSL, Boulder, Colorado, USA, from their website at https://psl.noaa.gov.

### Data analysis

All analyses were performed using the open-source statistical software programme R [29].

#### Tsetse Catches

Catches of tsetse per trap per day varied between months. We hypothesised that this was due in part to seasonal patterns in temperature and/or rainfall. Using the ‘glmmADMB’ package, we fitted a generalised linear mixed effects model (glmm) to the daily catches of male and female tsetse. We specified a negative binomial distribution for catches, with site and day of collection as random effects and month as a fixed effect.

Using the model outputs, we estimated the mean daily catches and their 95% confidence intervals for each month. To assess the impact of temperature and total rainfall, we also produced glmms of catch with monthly rainfall and/or temperature as explanatory variables. The significance of these variables was assessed by analyses of deviance (ANODEV) using the ‘Anova’ function. In addition to assessing relationships of monthly catch, and the simultaneous mean temperature and total rainfall, we also assessed whether rainfall and temperature from earlier months had an effect; previous studies have found relationships between abundance of tsetse and the mean saturation deficit, likely affected by rainfall, from earlier months [30].

#### Age structure

Previous studies of the age structure of tsetse have shown large and statistically significant inter-monthly variations in the distribution of Ovarian Categories (OC) [31]. To discover whether similar variation occurred with *G. f. fuscipes* in Uganda, we assessed the statistical difference in the monthly distributions of OCs for all consecutive months by Chi-squared tests. We also calculated the mean OC for each month and used Tukey’s multiple comparison test to assess the statistical significance of differences between means. To elucidate trends over a series of months, we carried out regression analyses to assess changes in the mean Ovarian Category and time (months) as a continuous function.

#### Adult mortality rates

We used a maximum likelihood method to estimate adult female mortality from the distribution of ovarian ages among samples of females captured in traps [32]. The derivation of the method is provided here in Supplementary Information file S1.

#### Infection rates

To assess the statistical significance of differences in the percentage of tsetse observed to be (i) infected with *Trypanosoma* as estimated by dissection and (ii) positive by PCR by logistical regression, we fitted data using a general linear model (glm) with a binomial error distribution and a logit link function, using method as a factor. The significance of explanatory factors was assessed by analyses of deviance (ANODEV) using the ‘Anova’ function.

For all analyses of catches, age structure and mortality rates, the 95% error limits are shown.

## Results

### Catches of tsetse

Between April 2013 and July 2014, the four traps caught a total of 12 512 tsetse across 238 trapping days. The mean daily catches of male and female tsetse peaked in November 2013 (Fig. 1A) and the lowest numbers were caught in April 2014; mean daily catches of males and females showed similar peaks and troughs, but the mean catch and range were greater for females (2.5-10.4) than males (0.7-2.7). Fitting a glmm to the data showed that Month (P<0.001, Deviance=678.08., df=1) and Sex (P<0.001, Deviance=69.96.14, df=15)] were statistically significant factors but there was no statistically significant interaction between them (P=0.515, Deviance=14.14, df=15). Catches of males and females were pooled for analyses of the relationships between abundance, monthly rainfall and mean monthly temperature. The results showed that there was no significant correlation between mean catch and the current temperature (P=0.282, Deviance=1.16, df=1) or rainfall (P=0.689, Deviance=0.16, df=1) but for rainfall only there was a significant relationship between catch and the total rainfall from earlier months, with the strongest correlation being between catch and the rainfall from three months earlier (P<0.01, Deviance=20.6, df=1). Analysis of the pooled data on sex showed that the predicted mean daily catch increased significantly with the rainfall from three months earlier (P<0.01, Deviance=19.72, df=1). Catches increased from 6.6 (3.40-12.84) tsetse/trap/day for dry (0 mm rain/month) months to 12.2 (5.96-24.98) tsetse/trap/day for the wettest (190.7 mm/month).

**Fig. 1.**
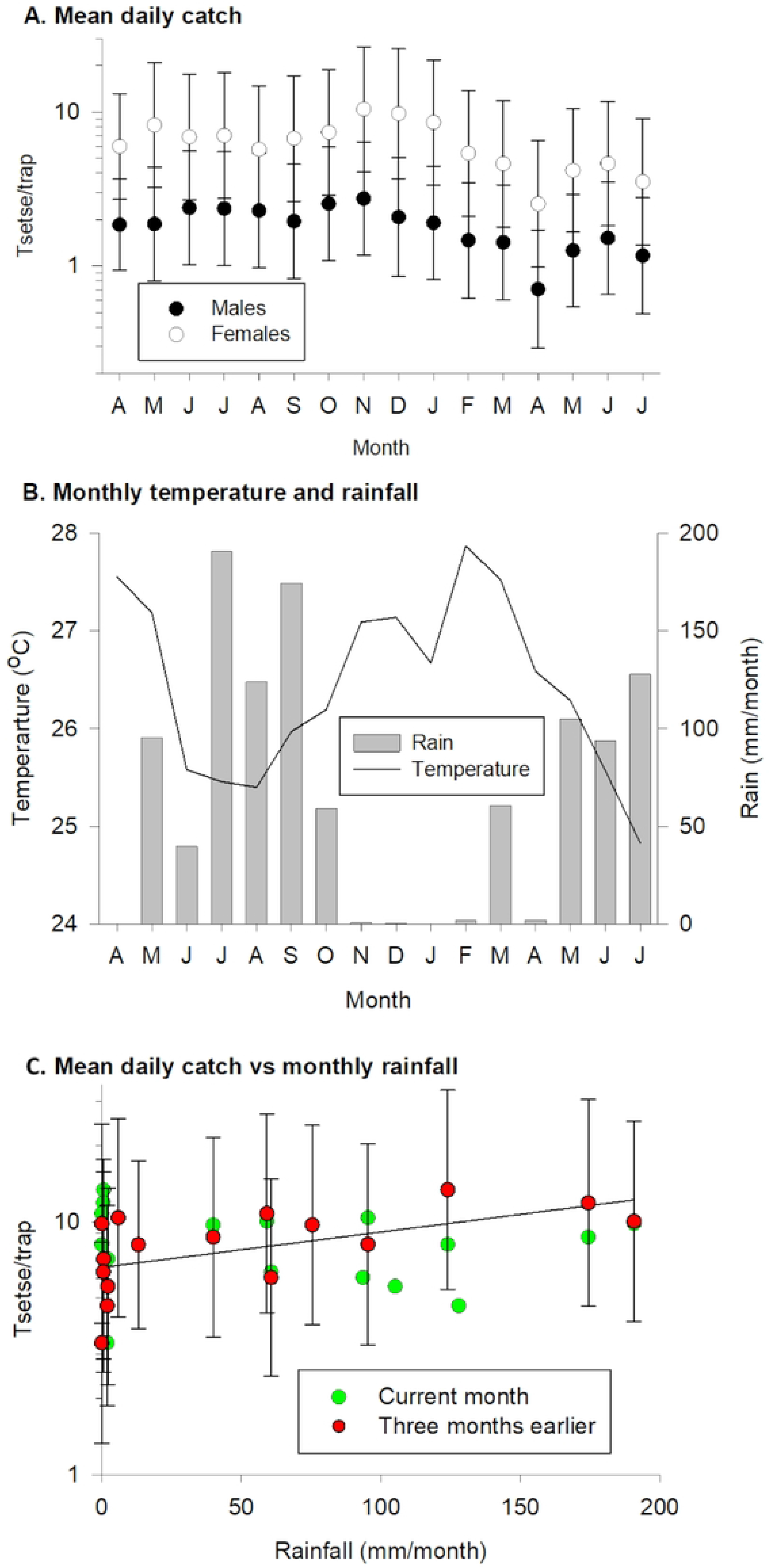
(A) Mean daily catch of male and female tsetse, (B) monthly mean temperature and total rainfall between April 2013 and July 2014 and (C) scatterplot of mean daily catch of tsetse (males and females combined) against monthly rainfall. For C, catches for each month are plotted against the rainfall for that month or that three months earlier. Line shows catch predicted from fitting glmm to rainfall three months earlier.

### Population age structure

A total of 5051 female tsetse were dissected to determine their Ovarian Category (Fig. 2). Comparing the distributions for all possible consecutive months showed that for most (12/15) contrasts there was no significant difference (Table 1). Two of the three significant contrasts occurred in the comparisons of May-June in 2013 and 2014 and the third was between October-November 2013. These changes were due largely to decreases in the percentage of tsetse from Ovarian Categories 5-7 and an increase in Ovarian Categories 0-3. The May-June changes in distribution were associated with a significant (Table 1) decrease in the mean Ovarian Category (Fig. 2 inset). The May-June period marks the transition between the hot-dry and wet seasons (Fig. 2B).

**Fig. 2.**
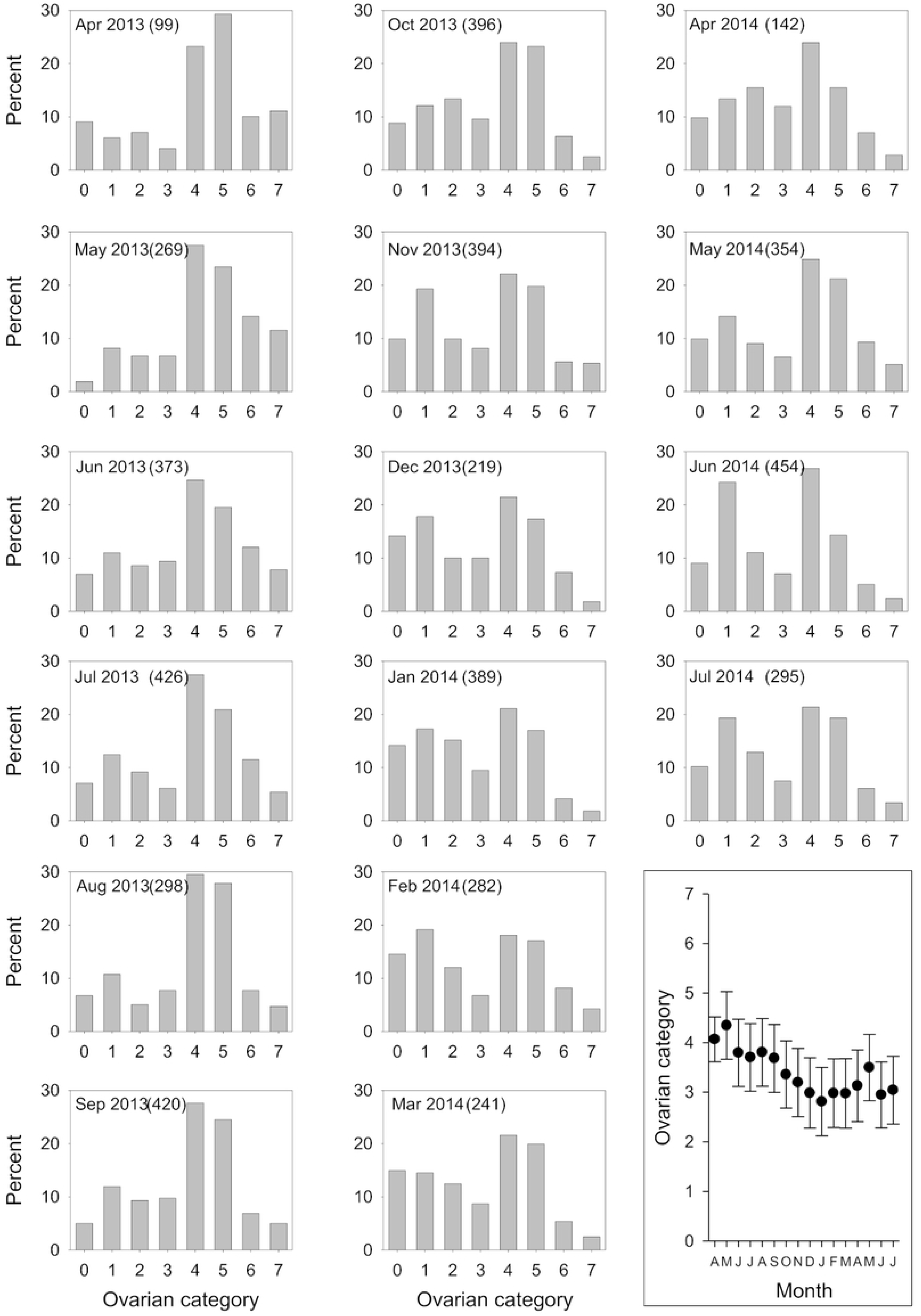
Percentage of tsetse in ovarian categories 0-7 for each month between April 2013 and July 2014. Inset shows mean ovarian category for each month.

**Table 1.**
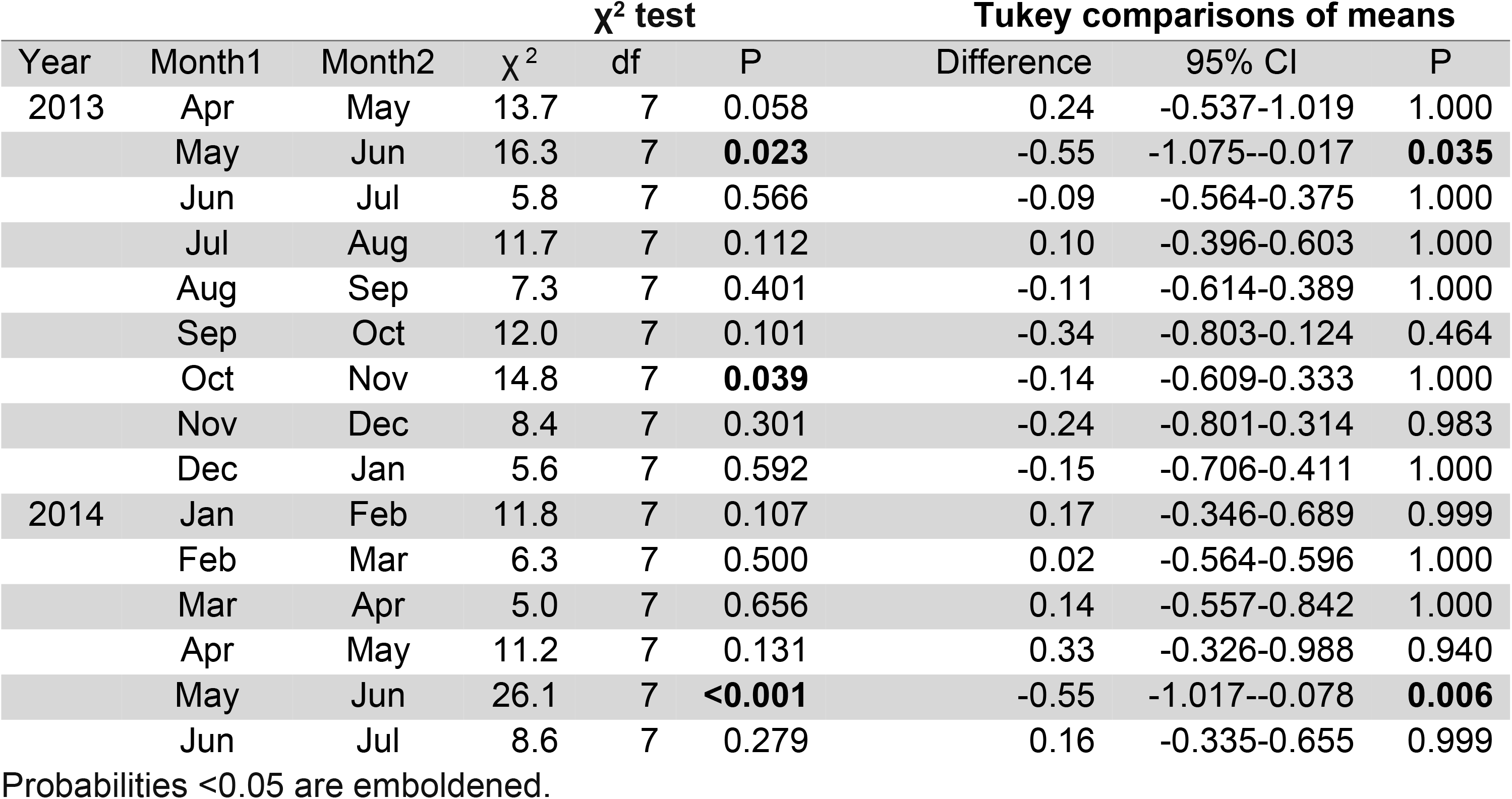
Statistical significance of differences in the distribution (χ^2^ test) and mean (Tukey test) of Ovarian Categories for consecutive months.

**Numbers in brackets show the sample size for each month.**

While the changes in age structure between consecutive months during the period June 2013-January 2014 were not statistically significant, there was a general and significant (P<0.001, Deviance=321.8, df=1) decline in mean OC (Slope=-0.146, SE=0.0156) during this period. Conversely, for the period January-May, where the OC distributions of consecutive months were also not significantly different, there was a statistically significant trend over the period (P<0.001, Deviance=81.1, df=1) with mean OC increasing (slope=0.155, SE=0.0343) between January-May 2014. Overall, mean OC declined during the wet season and increased during the dry, but the inter-monthly variation was generally small.

### Estimate of adult mortality

A global estimate of adult mortality, for the period June 2013 - May 2014, was calculated using the maximum likelihood (ML) method described in Supplementary File S1. The numbers of flies in ovarian categories 0 to 7+4n were 383, 564, 416. 334, 949, 815, 304 and 169, respectively, providing a mean mortality estimate for the 12-month period of 1.75% (1.63-1.88).

### Trypanosomes infection rates

A total of 1221 male and 5062 female tsetse were dissected and screened for trypanosomes using traditional microscopy methods. For males, trypanosomes were observed in 1.06% (13/1221, 0.57-1.81) of midguts, 0.08% (1/1221, 0.00-0.46) of salivary glands and 0.57% (7/1221, 0.23-1.18) of mouthparts. For females, the percentages were 1.50% (76/5062, 1.19-1.88), 0.12% (6/5062, 0.04-0.26) and 1.46% (74/5062, 1.15-1.83), respectively.

A sub-sample of 2369 tsetse (438 males, 1931 females) examined by microscopy between September 2013 and February 2014, were also screened using the mITS primers to identify tsetse positive for *T. brucei* s.l., *T. congolense* and *T. vivax* DNA. The results (Table 2) show that, for the midgut, salivary glands and mouthparts, the PCR-based method detected the presence of trypanosomes at a higher rate than microscopy. Over all combinations of sex and tissue, the numbers of positive tsetse detected by PCR were 1.8-9.3 times greater with PCR and the increases were statistically significant for all combinations apart from male mouthparts.

**Table 2.** Percentages of male (n=438) or female (n=1931) tsetse with trypanosomes detected by microscopy or PCR in the midgut (MG), salivary glands (SG) or mouthparts (MP), and the probabilities (P) that the percentages estimated by microscopy and PCR are not different (ANODEV).

| Tissue | Sex | n | Microscopy |  | PCR |  | F | df | P |
| --- | --- | --- | --- | --- | --- | --- | --- | --- | --- |
|  |  |  | mean | 95% CI | mean | 95% CI |  |  |  |
| MG | male | 438 | **0.68** | 0.141-1.989 | **3.65** | 2.102-5.865 | 10.0 | 1 | 0.002 |
|  | female | 1931 | **1.40** | 0.923-2.028 | **3.83** | 3.021-4.787 | 23.2 | 1 | <0.001 |
| SG | male | 438 | **0.23** | 0.006-1.265 | **1.83** | 0.792-3.567 | 6.3 | 1 | 0.012 |
|  | female | 1931 | **0.16** | 0.032-0.453 | **1.45** | 0.966-2.089 | 23.4 | 1 | <0.001 |
| MP | male | 438 | **0.46** | 0.055-1.640 | **1.14** | 0.372-2.644 | 1.3 | 1 | 0.247 |
|  | female | 1931 | **1.55** | 1.051-2.210 | **2.85** | 2.153-3.691 | 7.6 | 1 | 0.006 |

The three female tsetse that had infected salivary glands detected by microscopy, and hence nominally infected with *T. brucei*, were in ovarian categories 4-6; none of the salivary glands of 890 females in ovarian categories 0-3 were observed by microscopy to be infected. On the other hand, PCR detected trypanosome DNA in 2 females from each of categories 0-2 and 3 females in category 3 (SI, Table 1). However, only two of the tsetse in ovarian category 3 were identified as being from *T. brucei;* the other seven specimens were identified as being *T. vivax* or *T. congolense*.

Comparing the identity of infections based on the presence or absence of trypanosomes in the mouthparts, midgut and salivary glands and PCR showed (Table 3) that of the 23 tsetse identified as being *T. vivax* by microscopy, 17 were positive by PCR and all of these were indeed infected with *T. vivax*, albeit four were mixed infections with either *T. brucei* or *T. congolense*. Six tsetse putatively identified, by microscopy, as being infected with *T. congolense* were PCR-positive for T. congolense (n=4) or mixed infections with *T. brucei* (n=2). Finally, the three tsetse nominally infected with *T. brucei* were identified by PCR as two mixed positives (*T. brucei* and *T. congolense*) and one *T. congolense* positive. Tsetse observed with infections in the midgut only (n=20) were expected to have immature infections of *T. congolense* and/or *T. brucei*. Of these 20, 12 were successfully identified by PCR; 11 had one or both these species of *Trypanosoma*; and one was identified as *T. vivax*. In general therefore, the microscope- and PCR-based methods of identifying species of trypanosome infections were in broad agreement, but the PCR-based method was able to identify mixed infections.

PCR-based identity of tsetse (n) putatively identified by microscopy.

Table 3. PCR-based identity of tsetse (n) putatively identified by microscopy.
| Species | MP | MG | SG | PCR-based identity |  |  |  |  |  |  |  |
| --- | --- | --- | --- | --- | --- | --- | --- | --- | --- | --- | --- |
|  |  |  |  | n | None | Tv | Tc | Tb | Tv+Tb | Tv+Tc | Tb+Tc |
| *T. vivax* (Tv) |  |  |  | 23 | 6 | 13 | 0 | 0 | 2 | 2 | 0 |
| *T. congolense* (Tc) |  |  |  | 6 | 0 | 0 | 4 | 0 | 0 | 0 | 2 |
| *T. brucei* (Tb) |  |  |  | 3 | 0 | 0 | 1 | 0 | 0 | 0 | 2 |
| Tb and/or Tc |  |  |  | 20 | 8 | 1 | 5 | 3 | 0 | 0 | 3 |

Classically, the putative identity of infections is based on observation of trypanosomes in the mouthparts (MP), midgut (MG) and salivary glands (SG) as indicated by the black shading in the table.

PCR-based analyses of infection can detect single and mixed infections of *T. vivax* (Tv), *T. congolense* and *T. brucei*. None indicates tsetse observed to be infected by microscopy but not by PCR.

For trypanosomes detected by PCR, the majority of positives were located where expected:

*T. vivax* and/or *T. congolense* were detected in 83% (50/60) of PCR-positive mouthparts, *T. congolense* and/or *T. brucei* were detected in 68% (61/90) of PCR-positive midguts and *T*. *brucei* was detected in 14/36 PCR-positive salivary glands. However, we found many examples of species of *Trypanosoma* in unexpected places. For instance, *T. vivax* was detected in the midguts of 34 tsetse and *T. congolense* was detected in the salivary glands of 18 tsetse. These aberrant locations may be due to contamination during the dissection and processing of specimens. Evidence for this is provided by the observation that, of the 18 *T. congolense* positive salivary glands, only three occurred as a single tissue positive. For the other 15, *T. congolense* was detected in the mouthparts and/or midgut, where *T. congolense* is expected, as well as the salivary glands. In other cases, the detection of *Trypanosoma* species in unexpected tissues may be a consequence of ‘natural’ contamination of trypanosome DNA within the fly. For instance, it is not inconceivable that *T. vivax* in the mouthparts may become dislodged and ingested into the midgut. The 34 tsetse with midguts that were positive for *T. vivax* also had mouthparts that were positive for *T. vivax*.

Overall, and ignoring the locations of trypanosomes within the flies, the infection rates estimated by PCR-based analyses were 1.64% (1.321-2.244) for *T. brucei*, 2.41% (1.827-3.106) for *T. congolense* and 1.98% (1.461-2.630) for *T. vivax*.

### Blood meals

A subsample of 768 tsetse was analysed for the source of their bloodmeals; of these 131 bloodmeals (17.1%, 14.46-19.91) were identified successfully. The results (Fig.3) showed that cattle were the most common host (39%, 30.5-47.8) closely followed by humans (37%, 28.4-45.6). We believe that this is the first recording of Forest cobra (*Naja melanoleuca*) being identified as a host of tsetse (4%, 1.3-8.7).

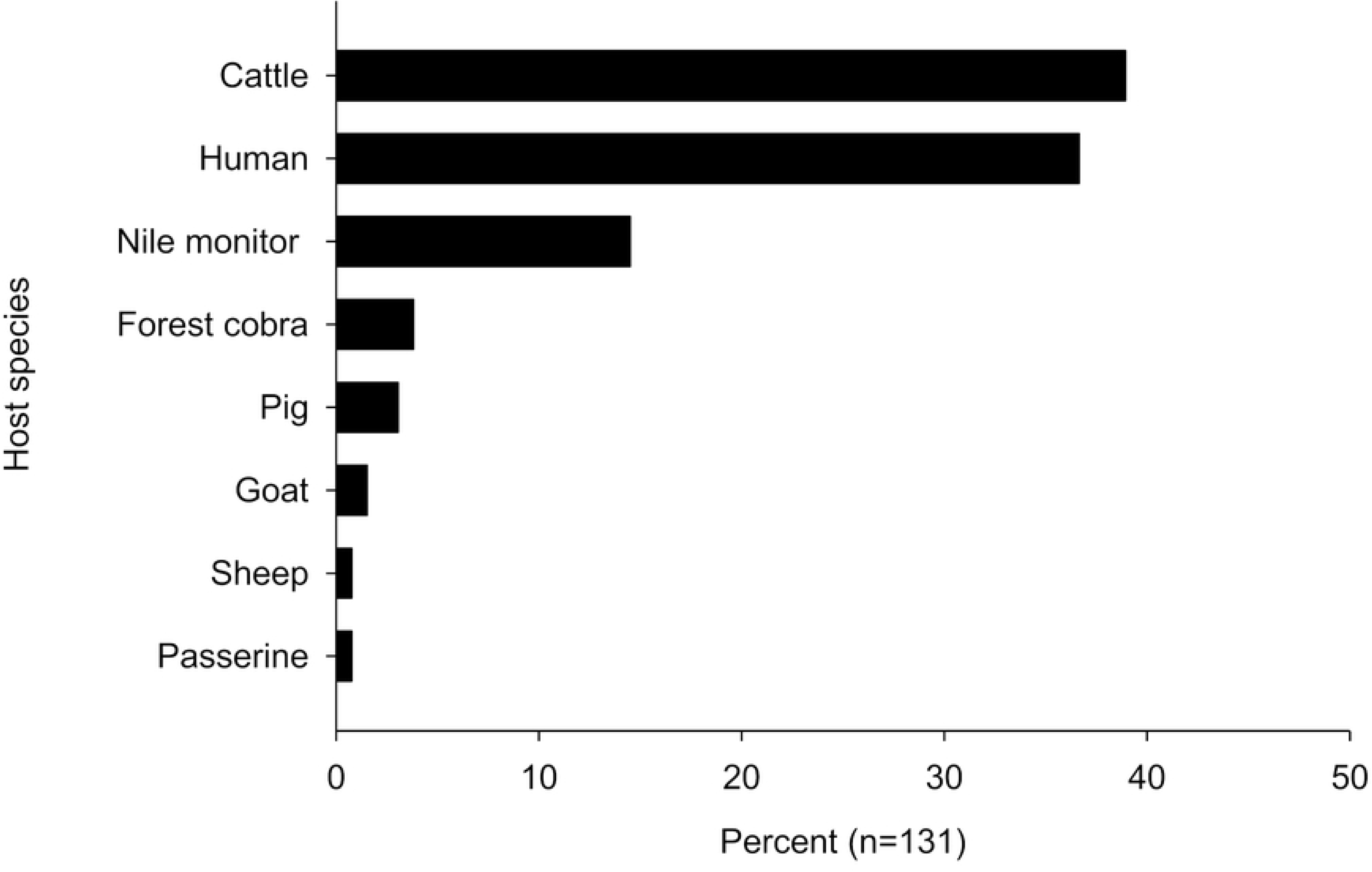

## Discussion

### Infection rates

By comparing estimates of infection prevalence estimated by microscopic examination of tsetse, and by a PCR-based method, we tested the null hypothesis that, for *G. f. fuscipes*, the two methods provide identical estimates of *Trypanosoma* infection prevalence. We reject this hypothesis since our results show that the latter was significantly more sensitive, particularly for infections of the midgut and salivary glands. Over all combinations of sex and tissues, the number of nominally infected flies were 1.8-9.3 times greater with PCR. Our results accord with studies of other species of savanna and riverine tsetse in, for example, Tanzania [14] and Cote d’Ivoire [15] respectively, but extend the findings to another species of tsetse for all the major pathogenic species of *Trypanosoma*. Assuming that the PCR-based method provides the more reliable estimate of infection rates, we estimate that, for *T. brucei*, *T. congolense* and *T. vivax* the rates were 1.64%, 2.41% and 1.98%, respectively.

While PCR-based methods suggested a higher rate of infection, the presence of DNA from a *Trypanosoma* species does not necessarily mean that the fly is infected; DNA from trypanosomes that have been killed by immune responses of the fly can be detected for a period after their ingestion [21]. Moreover, the presence of *T. brucei* or *T. congolense* in the midgut does not indicate an infectious tsetse. However, even if we assume that the detection of DNA always denotes the presence of viable *T. brucei*, then we detected only 2/890 (0.22%) female tsetse with mature salivary gland infections of *T. b. brucei.* This low rate of infection accords with previous findings, showing a low prevalence of salivary gland positives, equating to ∼1/1000 flies having a mature *T. brucei* sl infection [26, 33].

### Population dynamics

In collecting and analysing tsetse for infection with trypanosomes, we also gained insights into the population dynamics of *G. fuscipes* in north-west Uganda. First, the abundance of tsetse ranged between a maximum of 13.4 tsetse/trap in November 2013 and a minimum of 3.3 tsetse/trap in April 2014. There was no statistically significant variation in the ratio of males:females. This four-fold variation in catches contrasts with the much greater fluctuations in the abundance of savanna tsetse. In Zimbabwe for instance. catches of male *G. m. morsitans* and *G. pallidipes* in mopane woodland showed 35- and 13-fold differences between seasons respectively [34]. For females, the ratios were 2.5 and 7.5 respectively. Hence for these savanna species there is a large seasonal variation in numbers and also the ratio of males:females. Studies of other populations of savanna tsetse in East Africa also show large (>10-fold) variations in abundance [35, 36]. By contrast, and in accord with our results, natural variation in the abundance of riverine species of tsetse in East Africa seem to be much smaller [24, 37, 38]

While the fluctuations in catches were relatively slight, the change appeared to be correlated with rainfall 2-3 months earlier. Other studies have also found that rainfall [39], temperature [40] or saturation deficit and NDVI from earlier, rather than contemporaneous, months were correlated with the abundance or mortality rates of tsetse populations.

In keeping with the relatively stable numbers of tsetse, but contrasting with results for savanna tsetse [31], we found only slight seasonal variation in age structure. The only marked inter-month fluctuation seems to be associated with the onset of the rains at the end of the wet season (May-June). A potential explanation for the observed increase in the proportion of young tsetse at the beginning of the rains may be because pupae and/or teneral tsetse are susceptible to the dry conditions during the hot season (Dec-April). The survival of the pupae of *G. fuscipes* is reduced under dry conditions in comparison to savanna species of tsetse [41]. Poor survival of pupae and teneral tsetse may lead to an increase in the proportion of older tsetse in the population. The change in age structure might be greater than indicated by trap catches, since traps tend to catch only old flies [17].

Changes in pupal survival and humidity may explain the relationship between abundance of tsetse and rainfall 2-3 months earlier. There is a lag between the onset or cessation of the rains and changes in relative humidity [39]. As more humid conditions become established, there is increased survival of pupae leading to an increase in the numbers of adults about a month later, i.e. the duration of the pupal period. Catches of tsetse from traps are biased towards older flies; the most numerous age class for most months was Ovarian Category 4 (Fig. 2) where flies would be >40 days old.

The female adult mortality rate of 1.75% per day, estimated using the ML method applied to samples of flies collected over a 12-month period, is markedly lower than estimates averaging about 2.4% per day, resulting from the analysis of female *G. pallidipes* from Zimbabwe [31]. This is to be expected, given the harsher climatic conditions in the Zambezi Valley. Analysis of the Zimbabwe data showed that application of the method to samples of flies collected over monthly periods produced estimates that made little sense, given meteorological conditions at the times of capture. For example, mortality was found to take a minimum in December, one of the hottest months of the year; this is contrary to mark-recapture estimates indicating that mortality rates increase significantly with increasing temperature. This problem has been ascribed, primarily, to the fact that the estimation of mortality from age distribution involves the assumption of an underlying stable age distribution. As pointed out previously [42], it seems unlikely, given the large oscillations in temperature, in the Zambezi Valley, that these distribution are ever stable. Given the more equable climate of Uganda it seemed possible that there might be scope for estimating mortality over periods of time of order one month. In practice, however, it was found that the mortality estimates, as with the Zimbabwe situation, showed no sensible correlation with meteorological conditions.

### Bloodmeals

Our results show that cattle and humans were the most important sources of bloodmeals. The range of hosts is similar to results from earlier studies [43] but the proportion of meals from cattle and humans is higher than previously reported for *G. fuscipes* from this area of Uganda.

The high proportion of meals from humans combined with the low mortality rate is important because these factors contribute to higher rates of transmission. Building on Roger’s models of trypanosomiasis [44], many HAT models assume that natural tsetse mortality rates are 3%/day and the proportion of meals from humans is 30%. Substituting values of 1.75% and 37%, respectively, in Roger’s model increases R_0_ from 2.7, 388.2 and 64.4 for *T. brucei*, *T. congolense* and *T. vivax* (Table 3 in [44]) to 3.06, 610.8 and 114.9, respectively. The high proportion of meals from cattle suggests that regular treatment of cattle with pyrethroids would be an effective method of controlling tsetse in this area.

### Limitations

#### Estimates of infection rates

We detected trypanosome DNA in tissues where they are not expected: *T. vivax* was detected in the midgut of 34 tsetse and *T. congolense* in the salivary glands. At least some of these aberrant locations appear to be due to contamination by the dissectors despite our efforts to avoid this. The contamination was between tissues of the same fly rather than between flies and our procedures were less effective at preventing the former. While there is evidence of some contamination affecting our estimates, there is also evidence that detection of trypanosome DNA in unexpected tissues is also a natural phenomenon. For instance, a relatively high proportion of midguts with *T. vivax* were associated with the detection of *T. vivax* in the mouthparts. The simplest explanation of this is the passage of *T. vivax* parasites, or DNA, from an infected proboscis into the midgut during feeding and the subsequent detection of this transient DNA by PCR.

#### Seasonal variations in abundance and age structure

The study was conducted over 15 months in a relatively small geographical area; we use the results to infer large-scale seasonal patterns in populations of *G. f. fuscipes,* and the underlying causes of these changes. We are not aware of previous studies of this or longer duration which analysed abundance, age structure and diet. Nonetheless, we recognise that stronger inferences could be produced by conducting the studies over several years and over a wider geographical area. Data being produced by vector control operations conducted in north-west Uganda [37] will produce longer series over larger areas and future analyses of these data will allow us to build on the present results.

#### Estimates of mortality rates

The age composition of female tsetse shows some unexpected features, most noticeably the very large proportions of flies in ovarian category 4+4*n* and 5+4*n*.These are composite categories, which include flies that have ovulated 4, 8, 12,…etc and 5, 9, 13 …. etc times, and it is thus unsurprising that they contain greater proportions than found in category 3. However, since we expect numbers to decline monotonically with age, the differences between the proportions of flies in category 4+4*n*, or 5+4*n*, and those in category 3, should be less than the proportion in category 7+4*n*. In fact, however, the reverse is true for every one of the 16 months of the study – as is obvious from Figure 2.

These results are consistent with suggestions that there is a marked sampling bias against young tsetse (<Ovarian Category 4), as has been reported for savanna tsetse [17]. Such biases undermine confidence in our approach to estimating mortality rates. Research on the causes and consequence of biases in sampling methods are required.

### Conclusion

Our study has shown that PCR-based methods provide a more sensitive way of detecting and quantifying infection of tsetse with *T. congolense*, *T. brucei* s.l. and *T. vivax*. During the 15-month period of our study, there were only slight changes in the abundance, age structure and infection rates of tsetse. The changes that were observed were correlated with rainfall in previous months, which we suggest affected the survival of pupae.

The high proportion of meals from cattle suggest that insecticide-treated cattle should make a significant contribution to the control of HAT and AAT.

---

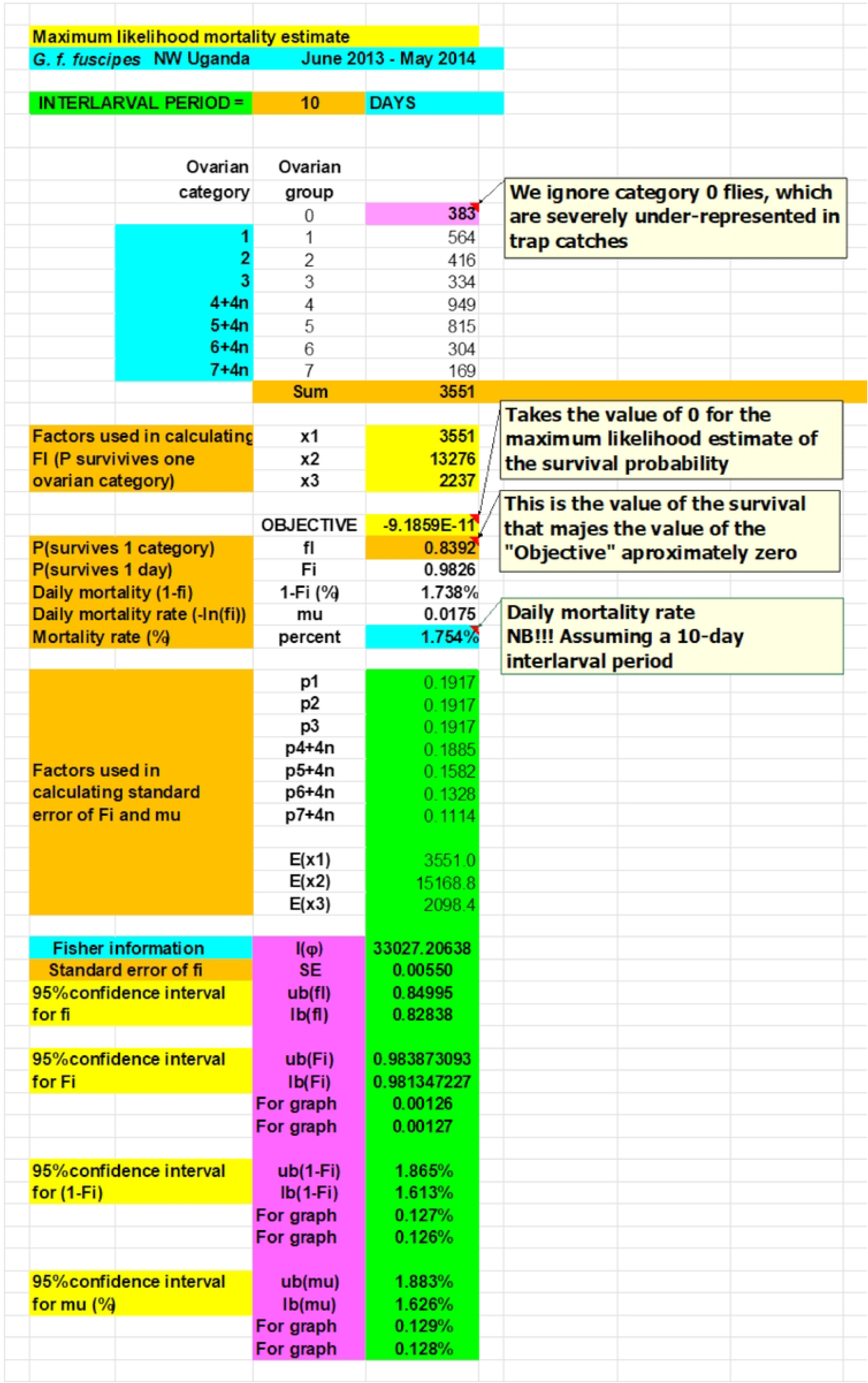

